# Comparative metagenomic analysis of biosynthetic diversity across sponge microbiomes highlights metabolic novelty, conservation and diversification

**DOI:** 10.1101/2022.04.01.486688

**Authors:** Catarina Loureiro, Anastasia Galani, Asimenia Gavriilidou, Maryam Chaib de Mares, John van der Oost, Marnix H. Medema, Detmer Sipkema

**Affiliations:** Laboratory of Microbiology, Wageningen University, Stippeneng 4, 6708WE Wageningen, The Netherlands; Groningen Institute for Evolutionary Life Sciences (GELIFES), University of Groningen, Nijenborgh 7, 9747 AG, Groningen, The Netherlands; Bioinformatics Group, Wageningen University, Droevendaalsesteeg 1, 6708PB Wageningen, the Netherlands

**Author notes:** Corresponding authors: Detmer Sipkema & Marnix Medema. Competing Interests MHM is a co-founder of Design Pharmaceuticals and a member of the scientific advisory board of Hexagon Bio.

## Abstract

Marine sponges and their microbial symbiotic communities are rich sources of diverse natural products (NPs) that often display biological activity, yet little is known about their global distribution landscape and the symbionts that produce them. As the majority of sponge symbionts remain uncultured, it is a challenge to characterize their NP biosynthetic pathways, assess their prevalence within the holobiont and measure their diversity across sponge taxa and environments. Here, we explore the microbial biosynthetic landscapes of three sponge species from the Atlantic Ocean and the Mediterranean Sea. This dataset reveals striking novelty in its encoded biosynthetic potential, with less than 1% of the recovered gene cluster families (GCF) showing similarity to any characterized biosynthetic gene cluster (BGC). When zooming in on the microbial communities of each sponge, we observed higher variability of both secondary metabolic and taxonomic profiles between sponge species than within species. Nonetheless, we also identified conservation of GCFs, with 20% of sponge GCFs being shared between at least two sponge species, and a true GCF core comprised of 6% of GCFs shared across all species. Within this functional core, we identified a set of widespread and diverse GCFs encoding nonribosomal peptide synthetases (NRPS) that are potentially involved in the production of diversified ether lipids, as well as GCFs putatively encoding the production of highly modified proteusins. The present work contributes to the small, yet growing body of data characterizing NP landscapes of marine sponge symbionts, and to the cryptic biosynthetic potential contained in this environmental niche.

## Introduction

Marine sponges (Porifera) are benthic heterotrophic filter feeders that harbour diverse and abundant communities of microbial symbionts in their tissues (1,2). These communities have aided marine sponges in their expansion across diverse ecological niches (2–4), and are dominated by Proteobacteria, Acidobacteria, Chloroflexi, as well as the sponge-specific Poribacteria (5–7). The complex unit of a sponge and its microbial consortium is referred to as the ‘holobiont’(8), and can be divided into two categories based on the abundance and diversity of microbes in the sponge tissue, with high microbial abundance (HMA) sponges harbouring richer and more diverse communities (7,9–11), at 10^8^ –10^10^ microbial cells g^−1^ sponge wet weight (5,7), and low microbial abundance (LMA) sponges hosting on average 10^5^ –10^6^ microbes g^−1^ sponge wet weight(9). Although there are exceptions to this rule (12), there is commonly an enrichment of Poribacteria, Chloroflexi and Acidobacteria in HMA sponges (11). Additionally, there appears to be a functional microbial core which displays gene abundance differences in core metabolic functions (13,14).

Natural Products (NPs) are ubiquitous small molecules that play a key role in symbiosis, as mediators of interactions within the holobiont (15,16). In this generally mutualistic relationship, the microbes provide their host with chemical compounds to prevent predation, fouling and infection, while receiving primary metabolic nutrition and a hospitable habitat (17–20). Whilst symbiosis often leads to the reduction and specialization of a symbionts’ genome, the sponge holobiont secondary metabolism is maintained through positive selective pressure (21,22). This specialization can also lead to the generation of ‘super producers’ with high numbers of biosynthetic gene clusters, such as the genus “*Candidatus* Enthotheonella”, that was first discovered in sponges (15,23). Sponge symbionts are a particularly prolific source of diverse NPs that often display biological activity (3,24,25). However, the large majority of sponge symbionts remain uncultured, which has hindered the characterization of NP-mediated host-symbiont interactions (26,27), and, consequently, the access to this untapped reservoir of a broad spectrum of bacterial secondary metabolites (28,29).

The enzymes that catalyze the production of specific secondary metabolites are generally encoded in Biosynthetic Gene Clusters (BGCs) (30). BGCs discovered in marine symbiotic systems display a high degree of diversity, with non-canonical cluster architectures that allow for the biosynthesis of highly diverse secondary metabolites (23,28,31,32). Still, there is a relatively low number of studies examining the sponge holobiont with a focus on secondary metabolism (28,33,34). Metagenomics aids in recovering environmental genetic material from uncultured microbes, and reconstruction of metagenome-assembled genomes (MAGs) facilitates studying these gene clusters within their genomic and taxonomic context (35–37). Recent development of tools like antiSMASH (38) and BiG-SCAPE (39) (the most widely used tools for prediction and comparison of BGCs) as well as MIBiG (30) (a curated database for BGCs with experimentally determined products) allow for efficient leveraging of metagenomic sequencing datasets for discovery of BGCs. This approach has revealed the existence of uncharacterized microbes with diverse natural product repertoires (40,41), and facilitated linking putative secondary metabolites to their bacterial producers (29,31,42).

Untargeted mining of sponge microbiome metagenomes allows for a more complete view of the holobiont’s biosynthetic diversity and conservation across sponge holobionts (15). While NPs have mostly been described as highly niche-specialized metabolites, there is emerging data pointing to the conservation of certain BGCs in a symbiotic context (15,43). A broader view on the extent to which BGCs are shared between different sponge species, however, is still missing. Here we make use of these culture-independent methods to explore the combined taxonomic and biosynthetic landscape of marine sponge bacterial symbiont communities. In this way we establish a detailed overview of secondary metabolic diversity in sponges, which comprises a diverse array of mostly uncharacterized Gene Cluster Families (GCFs). The ubiquitousness of a part of this array supports the hypothesis of an ecologically important set of secondary metabolic pathways conserved among HMA sponge species. This includes a novel set of nonribosomal peptide synthetase (NRPS)-like GCFs associated with the production of molecules related to vinyl ether lipid phosphatidylethanolamine (VEPE) (44), as well as multiple GCFs spanning several ribosomally synthesized and post-translationally modified peptide (RiPP) classes that appear unique to sponge microbiota. In addition, we have identified the putative bacterial hosts of these BGCs, which constitutes an important next step towards accessing the biosynthetic potential that is still largely untapped in sponge microbiomes.

## Materials & Methods

### Sponge collection

*Geodia barretti* samples were collected and processed in (45). Atlantic Seawater (Seawater ATL) samples (gb1_f – gb10_f) were collected onboard R/V Hans Brattström of the University of Bergen from Korsfjord, Bergen, Norway (60°8.13’ N, 5°6.7’ E) in September and October 2017 by filtering 2 L of seawater through PVDF membrane filters (Merck Millipore, Burlington, Massachusetts,USA), pore size: 0.22um, diameter: 47mm. Filters were snap frozen in liquid nitrogen and stored at -80 °C. *Aplysina aerophoba* and Mediterranean Seawater (Seawater_MED) samples were collected and processed in (46), and are publicly available (46). *Petrosia ficiformis* sampling took place in August 2018 at a semi-submerged marine cave (5-6 m depth) with internal freshwater springs in Sfakia, Greece (35°12’ N, 24°7’ E) in a collaborative effort with the Hellenic Centre for Marine Research (HCMR). Immediately after collection, the samples were transferred to liquid nitrogen and stored at - 80 °C.

### Total DNA extraction and metagenomic sequencing

*A. aerophoba* (46), *G. barretti* and *P. ficiformis* sponge samples were crushed in liquid nitrogen to a fine power with pestle and mortar. Two hundred mg of sponge tissue powder was further disrupted by bead beating using milling balls (5×2 mm + 2×5 mm) and 2 steps of shaking for 20 s at 4,000 rpm in a Precellys 24 tissue homogenizer (Bertin Instruments, Montigny-le-Bretonneux, France) (47). Tissue lysate was further used for DNA extraction with the AllPrep DNA/RNA/Protein Mini Kit (Qiagen, Hilden, Germany). Total DNA extraction of seawater filters was done using the entire filter, following the protocol described above. DNA extracted from water filter replicates gb5_f and gb6_f was pooled prior to sequencing to meet minimum DNA quantity requirements. The extracted DNA was further cleaned by a collagenase treatment using C9891 Collagenase from *Clostridium histolyticum* (Sigma-Aldrich, St. Louis, Missouri, USA) at a concentration of 2.5 mg/mL for 30 min at 4 °C vortexing every 5 min at max speed for 10 s. It was then purified using the MasterPure Gram Positive DNA purification kit (Lucigen, Middleton, Wisconsin, USA) using the manufacturers’ instructions, and passing through Illustra MicroSpin S-400 HR columns (GE Healthcare, Chicago, Illinois, USA). *G. barretti, P. ficiformis* and Seawater_ATL total DNA was obtained, that was sequenced by Novogene (Hong Kong, China) using the Illumina HiSeq PE150 platform. *A. aerophoba* and Seawater_MED total DNA was sequenced by the Research Group Genome Analytics (GMAK) at DSMZ (Braunschweig, Germany) using Illumina HiSeq PE100. *A*.*aerophoba* DNA was also prepared for Pacific Biosciences (PacBio, Menlo Park, CA, USA) long-read sequencing and was processed as in (48) and sequenced at DSMZ. Complete sample metadata can be found in Supplement 1.

### Quality trimming and adapter removal

Illumina HiSeq read adapter removal, quality filtering and normalization was done using the BBduk.sh script from BBTools suite v37.64 (49), following user guide indications, with parameters ktrim=r k=23 mink=7 hdist=1 tpe tbo qtrim=rl trimq=20 ftm=5 maq=20. The minlen parameter was set to 30 for *A. aerophoba* and Med_SW samples, and to 50 for all remaining samples. BBDuk (49) was also used to remove sequencing artifacts and phi X contamination, with default settings.

### Metagenomic assembly

Reads were normalized for coverage with BBNorm (49) with parameters target=100 min=5 for *Petrosia ficiformis*, and target=200 min=3 for all other samples. As SPAdes v3.12 (50) hybrid mode (--pacbio) does not support co-assembly, *A. aerophoba* sample Aply22 sequencing replicates were merged prior to hybrid assembly. *A. aerophoba* filtered Illumina HiSeq reads and PacBio reads were assembled with SPAdes v3.12 (50) using the --meta and - -only-assembler flags. Filtered Illumina HiSeq reads from all other samples were assembled with SPAdes v3.12 (50) using the --meta and --only-assembler flags.

### MAG binning and classification

Contigs were binned using metaWRAP v1.2 (51) with minimum completeness of 75% and maximum contamination of 10%, using MaxBin2 (52), metaBAT2 (53), and CONCOCT (54). The obtained bins were dereplicated using dRep (55) v2.5.4 with default parameters for primary clustering and secondary clustering using parameters --S_algorithm gANI --S_ani 0.95 (56). The dereplicated bins were taxonomically classified using the GTDB-Toolkit (57) v1.1.0 (GTDB-Tk) classify workflow. A phylogenetic tree of the dereplicated bins was created using the multiple sequence alignment generated in the GTDB-Tk (57) align workflow with FastTree (58) v2.1.11, default parameters. This tree was visualized and annotated using the Interactive Tree of Life (59) (iTOL v6) online tool.

### Metagenome 16S rRNA gene taxonomic classification

16S rRNA gene sequences were extracted and characterized using Phyloflash (60) v3 with SILVA rRNA gene database (61) v38.1, parameters -taxlevel 6 -poscov and -readlength 150 for *G. barretti*, Seawater_Atl and *P. ficiformis* samples, 100 for *A. aerophoba* and Seawater_Med Taxa relative abundance plots (--task barplot –level 6) and a nearest taxonomic unit (NTU) table (--task ntu_table --level 6) were generated with phyloFlash_compare.pl.

### Biosynthetic gene cluster analysis

BGC prediction was performed for all contigs over 4000 bp in length using antiSMASH(38) v 5 using the following parameters: --cb-general --cb-subclusters --cb-knownclusters -- minlength 4000 --hmmdetection-strictness relaxed --genefinding-tool prodigal-m -- clusterhmmer --asf --smcog-trees --pfam2go. BiG-SCAPE (39) v1.0.1 was run on all predicted BGCs using parameters --mix -v --mode auto --mibig --cutoffs 0.5 --include_singletons. BiG-SCAPE (39) network files were processed by in-house Python scripts available at https://github.com/CatarinaCarolina/sponge_meta_BGC to generate Fig. 1a, b, e as well as Fig. 5 using the Python package UpSetPlot v0.4.1. Phylogenetic analysis of NRPS-like GCFs was conducted using CORASON (39) v1, default parameters, with gb8_2 contig 859 gene 9 as a query and MIBiG BGC0000871.1 as reference BGC. A representative BGC was selected from all GCFs captured by CORASON(39) and the respective AMP domain amino acid sequence was used in a multiple sequence alignment (MSA) done with Muscle (62) v3.8.31. FastTree (58) v2.1.11 processed the MSA into a phylogenetic tree, visualised with iTOL (59).

**Figure 1.**
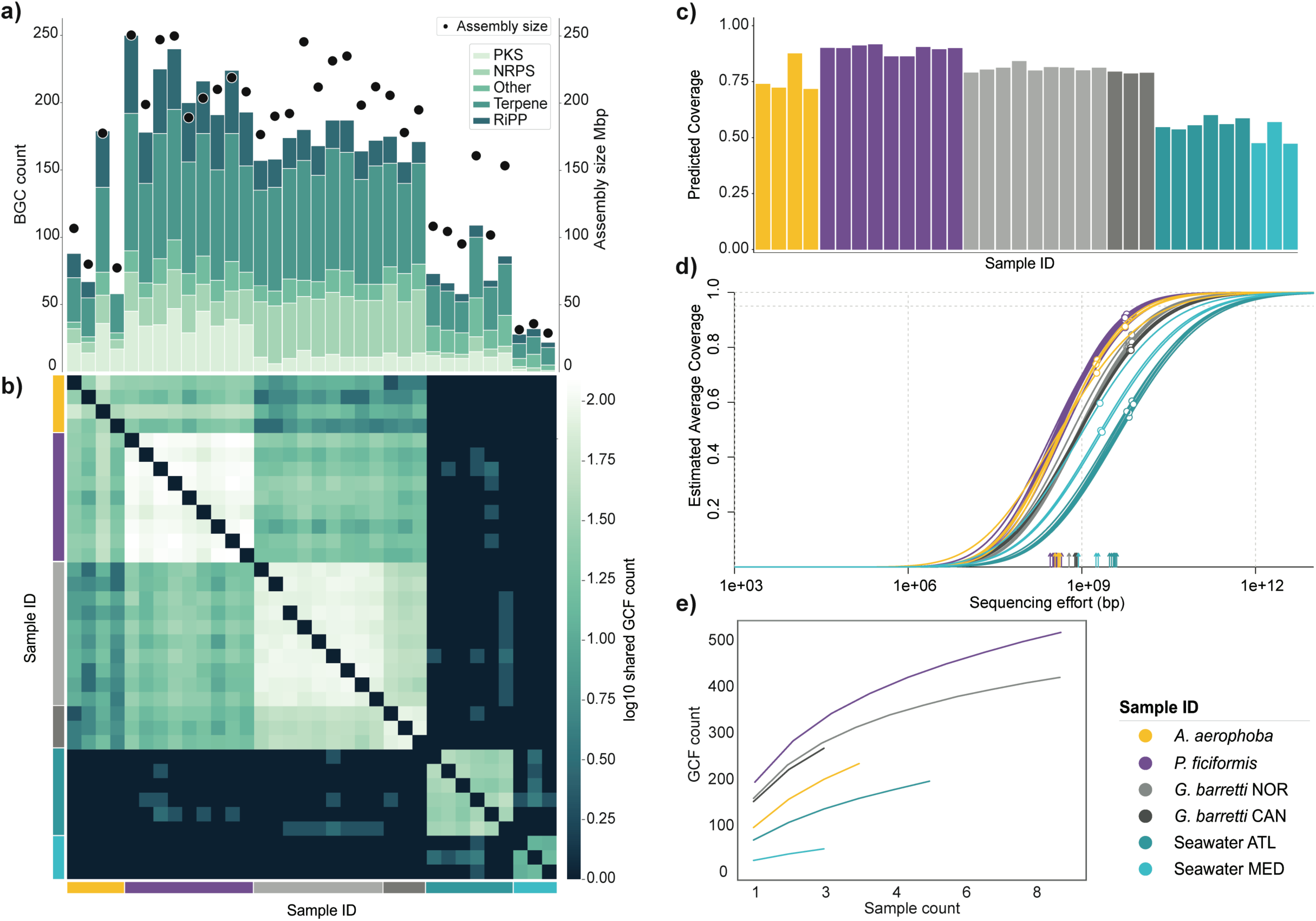
Biosynthetic gene clusters (BGCs) and gene cluster families (GCFs) in marine sponges. **a)** BGC counts coloured per major class (right y axis, stacked bars) and assembly size (left y axis, black dots) per sample. Assembly size includes only contigs above 4 000 bp length, used as input for antiSMASH; **b)** Log10-normalized pairwise heatmap of shared GCF counts between samples **c)** NonPareil sequence coverage estimates; **d)** NonPareil curves estimating the relationship between estimated coverage (y axis, white dots indicate sample estimated coverage) and sequencing effort (x axis, vertical arrows indicate sample sequencing effort); **e)** GCF rarefaction curves.

A RiPP-specific BiG-SCAPE (39) run was carried out by selecting all RiPP BGCs previously identified, excluding those classified with the antiSMASH (38) rules DUF692 and TIGR03975, as these BGCs were likely false positives based on manual inspection. GCF abundance, i.e. normalized RPKM (reads per kilobase per million), was calculated using BiG-MAP (63) with parameters -tg 0 -c 0.5 for the family module, and otherwise at default settings.

### Sequence estimated coverage

Sample read redundancy estimation was calculated using Nonpareil (64) v3.304 with parameters -T kmer -X 1000000. Nonpareil curves were built using the R package Nonpareil in RStudio, R v4.0.3.

### Statistical & diversity analysis

The Phyloflash (60) bacterial/16S-rRNA-gene-derived NTU table, normalized by relative abundance, and the BiG-MAP (63) normalised RPKM table were used to calculate Sample Shannon α-diversity scores with the Python package skbio.diversity.alpha.shannon, as well as to generate a Bray-Curtis dissimilarity matrix with the Python package scipy.spatial.distance.braycurtis. Pearson’s correlation score for Shannon α-diversity scores was calculated using the Python package scipy.stats.pearsonr. A Principal Coordinate Analysis (PCoA) using these same distance matrices was performed using the Python package skbio.stats.ordination.pcoa. Permutational Multivariate Analysis of Variance (PERMANOVA) was calculated for both distance matrices grouping by sample type using the Python package skbio.stats.distance.permanova.

### MAG & GCF data integration

MAG iTOL (59) annotation tables were generating by processing GTDB-Tk and BiG-SCAPE outputs with in-house Python scripts available at https://github.com/CatarinaCarolina/sponge_meta_BGC.

## Results & discussion

### Sponges harbor a striking number of novel biosynthetic gene clusters

IIn total, 5 082 biosynthetic gene clusters (BGCs) were detected in the sponge and seawater samples, which were grouped into 1 186 gene cluster families (GCFs) plus 394 singletons. Of all sponge-derived GCFs, only four included experimentally characterized reference BGCs from MIBiG (65), a striking display of the potential novelty contained within marine sponges. We can observe differences between sponge species with regard to their biosynthetic gene cluster (BGC) counts as well as sequencing depth and assembly size (Fig. 1a). The extended period between samplings and the diverse nature of the sequencing performed for each sample set warranted looking into a possible impact of the type of sequencing performed on sequence coverage and genomic content recovery. Although BGC count patterns and assembly size/sequencing effort do follow the Nonpareil estimated coverage, the latter shows less amplitude in variation (Fig. 1c, d). This indicates that BGC count patterns (Fig. 1a) observed are largely representative of the samples’ inherent sequence diversity.

As BGC boundaries are notoriously difficult to define and BGCs are often fragmented, we have from here onwards considered the GCF to be the biosynthetic unit of study, in order to minimise inflation originating from fragmented BGCs. Specialized metabolic similarity between samples can then be estimated based on the fraction of shared GCFs (Fig. 1b), with sponge samples showing high similarity even across species and habitat locations, but not to seawater samples (Fig. 1b). GCF discovery rarefaction curves (Fig. 1e) show that the biosynthetic diversity in HMA sponges is not fully captured and indicate that a higher sequencing effort would be needed for complete coverage. Nonpareil projection curves (Fig 1d) estimate the required sequencing effort at 10-100 Gbp for 90% sequence diversity recovery for HMA sponges. However, with estimated coverage values above 75% for all sponge samples, our data recovers a large fraction of this diversity, allowing characterization of the lion’s share of the biosynthetic landscape of these holobionts.

When investigating the distribution of GCFs across the dataset we see that each sponge species retains a large fraction of unique GCFs (65% of all sponge GCFs). Furthermore, there is also significant individuality between the two *G. barretti* sample groups, i.e. samples from two different geographical locations (Norway and Canada), with unique GCFs outnumbering shared GCFs in each of the geographical locations. Nevertheless, there is also a clear indication of specialized metabolite conservation across sponge species, with all sponge species sharing a core of 6% (58 GCFs) of all their encoded GCFs (Fig. 2). We recorded that an additional 200 GCFs were shared between at least 2 of the 3 sponge species. In the sponge holobiont, functional redundancy has previously been identified for primary metabolism, with nutritionally specialized guilds that span several taxonomic affiliations (66). With regard to specialized metabolism, conservation across species has been shown with the Sponge Ubiquitous Polyketides (SUP) cluster (67), the Sponge Widespread Fatty Acid Synthase (swf) cluster (68) and the Sponge derived RiPP Proteusins (srp) (15). Furthermore, Mohanty *et*.*al*. (69) recently reported that the presence of bromotyrosine alkaloids, signature NPs that are present across phylogenetically distant sponges, is not dependent on the sponge microbiome taxonomic architecture. We recovered both SUP gene clusters (67) and swf-like gene clusters (68) from all sponge species (Supplement 2.1), as well as srp-like clusters from two of the three species (discussed below, Fig. 7).

**Figure 2.**
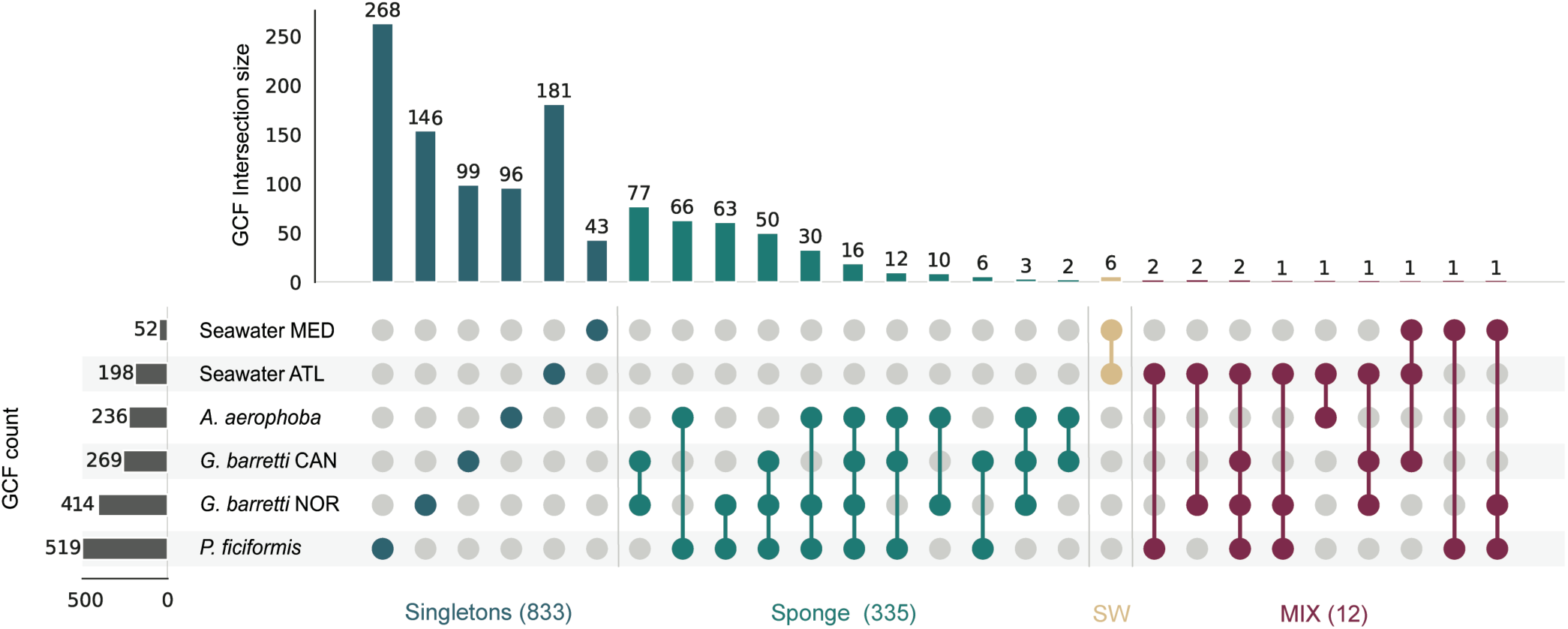
GCF intersections across sample groups. UpsetPlot illustrating GCF intersections between sample groups, i.e. samples grouped by sponge species and sampling geographical location. GCF count refers to the total count for each sample group, intersection size refers to the GCF count of each fraction/intersection (grid dots). For each category (singletons, sponge, mix of sponge and seawater) with >1 intersection the total GCF count is shown in brackets.

### Taxonomic diversity and Biosynthetic Gene Cluster Family diversity follow similar trends

As it remains unclear from which taxon of the holobiont the predicted BGCs and GCFs derive, we aimed to gain insight into the relations between functional gene content and taxonomic diversity. Observed prokaryotic community composition (Fig. 3a) follows the commonly observed profiles (70,71), with Proteobacteria, Chloroflexi, Acidobacteria and Poribacteria occupying the largest fractions of these sponge-associated bacterial communities.

**Figure 3.**
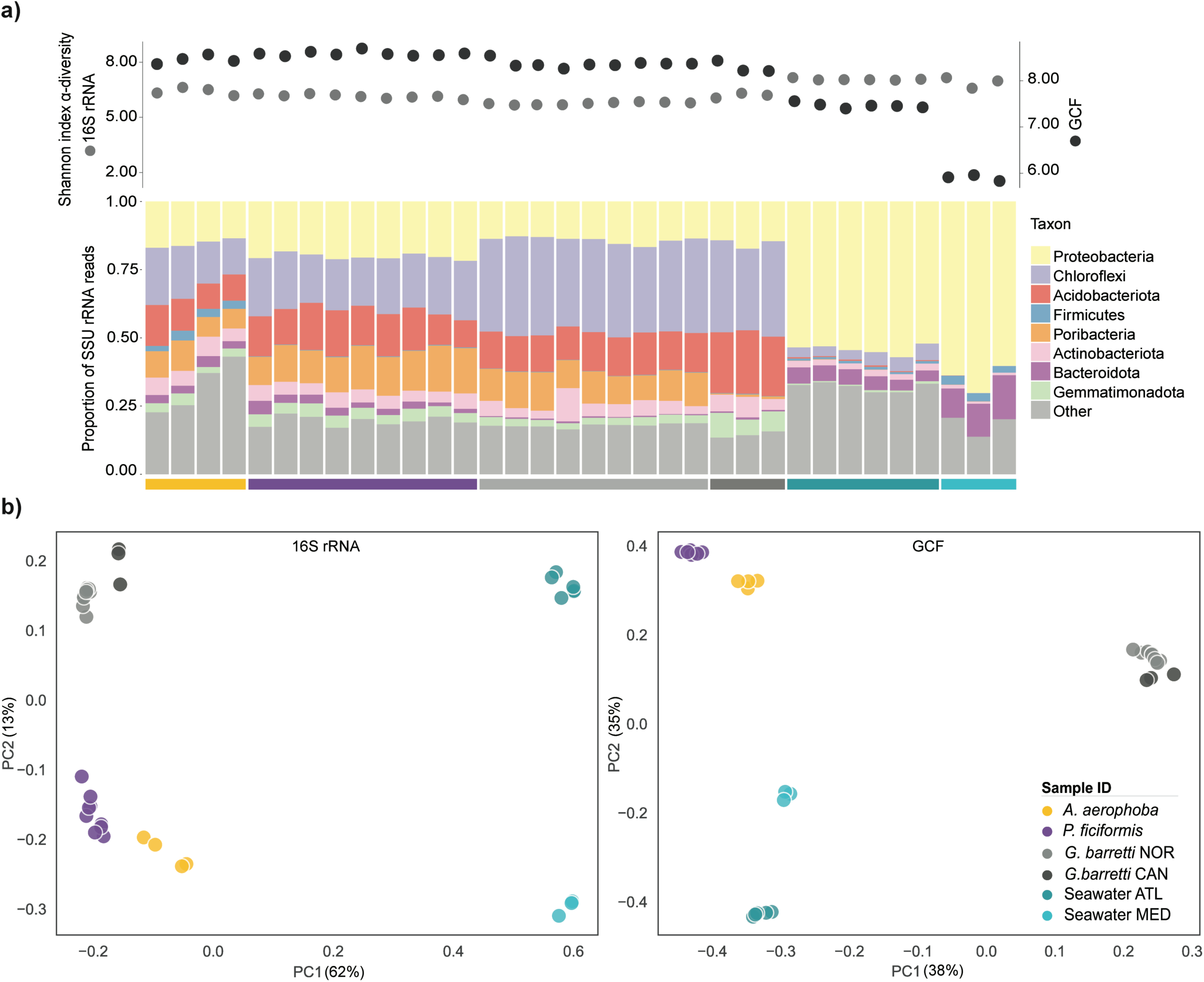
Biosynthetic potential and taxonomic diversity comparison of marine sponges. a) Sample Shannon index α-diversity scores based on 16S SSU rRNA genus level NTU (left axis, light gray) and GCF (right axis, dark grey) content, and sample prokaryotic taxonomic profile classified at phylum level. b) β-diversity of sponge samples based on 16S SSU rRNA genus level NTU (left) GCF (right) content. Percentage explained variance is denoted on each axis label.

**Figure 4.**
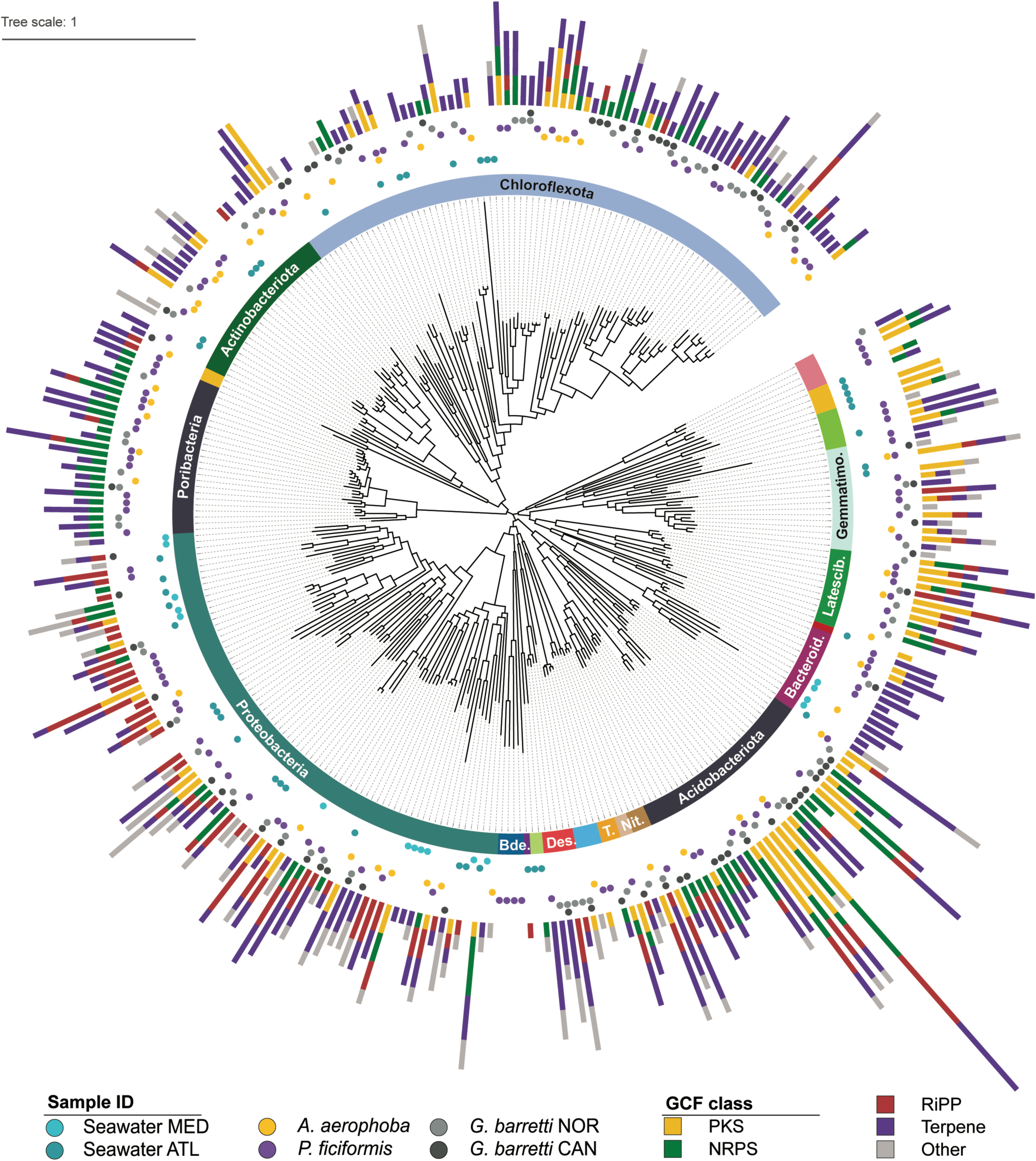
Gene Cluster Families present in the recovered MAGs. Cladogram based on GTDB classification of MAGs constructed in this study. Layer 1: phylum classification (truncated phyla are Gemmatimonadota, Latescibaterota, Bacteroidota, Nitrospinota, Nitrospinota_A Tectomicrobia, Desulfobacterota*_*B, and Bdellovibrionota). Complete MAG classification is in Supplement 4. Layer 2: Sample groups where MAGs are observed. Layer 3: GCF content coloured by class.

GCF-based Shannon α-diversity scores do not follow taxonomy-based α-diversity scores across the dataset (Fig. 3a), indicating that a higher genus diversity does not directly lead to a more diverse biosynthetic profile (Spearman’s r = -0.4, *p* = 0.01). Furthermore, we observe that there are significant differences between each sponge species’ taxonomy-based α-diversity scores (Kruskal-Wallis’ H = 17, *p* = 0.001), as well as GCF-based α-diversity scores (Kruskal-Wallis’ H = 19, *p* = 0.0001), with *A. aerophoba* showing the highest mean taxonomy-based α-diversity and *P. ficiformis* showing the highest mean GCF-based α-diversity (Supplement 3).

With respect to β-diversity of prokaryotic community composition and GCF composition, we observe that all sample groups are different, with taxonomy and GCF content generating similar patterns (Fig. 3b). The shallow Mediterranean species, *A. aerophoba* and *P. ficiformis* have relatively similar prokaryotic communities and GCF distributions that are different from those of the deep Atlantic sponge *G. barretti* from Norway and Canada. In addition, the prokaryotic communities and GCF distribution in seawater were different for Atlantic and Mediterannean seawater and also different than those of the three sponge species. PERMANOVA testing based on 16S rRNA gene genus-level NTU and GCF content revealed that all sample groups are significantly different (*p* = 0.001) in both contexts (Supplement 3). These results further support the notion that, despite the presence of a shared GCF core, each sponge species has a distinct encoded specialized metabolism profile.

### Acidobacteriota and Latescibacterota are the BGC power houses

We recovered a total of 313 metagenome-assembled genomes (MAGs), all classified at least at the phylum level (GTDB r95), from the sponge and seawater metagenomes (Fig.4, Supplement 4.2). We observe a relatively low sharedness of MAGs, with only 10% of bins being found in more than one sample group. This is in line with the recent work by Robbins *et al*. (33) who phylogenetically characterised 1 200 MAGs derived from 30 sponge species and reported similar patterns of shared vs. exclusive MAGs, with the presence of taxa that are unique to their host sponge species, as well as populations of Actinobacteriota and Acidobacteriota that are shared across sponge species. When the MAGs recovered here are shared, it is mostly so within the same sponge species (*G. barretti* NOR and CAN) or between sponges sharing similar habitats (*A. aerophoba* and *P. ficiformis*).

Of the 266 MAGs recovered from sponge samples, 96% contained GCFss, and of the 58 MAGs recovered from seawater samples 83% contained GCFs. Additionally, we observe an average of 3.5 GCFs per sponge MAG and 2 GCFs per seawater MAG. Acidobacteriota have been known as talented NP producers (40,72), and emerge also here as the most productive and functionally diverse phylum, represented by 28 MAGs with an average of 6.6 GCFs per MAG. Another diverse and consistently talented phylum is Latescibacterota (average 4.4 GCFs per MAG), comprising well-known members of sponge holobiont communities (28,33,34) that have only recently been linked to NP production (15), and thus constitute a potential untapped hub for discovery. We observe that members of both these phyla are present in all three sponge species. Additionally, two other prolific phyla, yet represented by fewer MAGS, are Nitrospirota (average 8.3 GCFs per MAG) and Desulfobacterota*_*B (average 6.0 GCFs per MAG). The candidate phylum Tectomicrobia is also represented here by three MAGs, respectively encoding for 2, 5, and 7 GCFs, and were classified as genus ‘SXND01’ within the family *Entotheonellaceae. Ca*. Tectomicrobia is famous for its biosynthetically prolific candidate genus *Entotheonella* (23), with other lineages within *Ca*. Tectomicrobia currently described as biosynthetically poor (45).

The previously mentioned SUP (67) and swf-like (68) GCFs that were obtained from all sponge species appear to be independent of symbiont taxonomy. They were found in several MAGs of the phyla Chloroflexota, Spirochaetota, Proteobacteria and Acidobacteriota in the case of SUP-like GCFs, and Nitrospirota, Latescibacteriota and Acidobacteriota in the case of swf-like GCFs. Multiple MAGs encoding SUP and swf*-*like GCFs belonging to different phyla were observed within the same sponge holobiont. However despite their similarity at the phylum level, at the species level (<95% gANI) the majority of these MAGs are specific to a single sponge species (Supplement 2.2).

### Widespread ether-lipid-associated BGCs in sponges

Within the shared GCF core, we identified nine Non-Ribosomal Peptide Synthetase (NRPS)-like GCFs with a similar architecture: a NRPS-like core gene (*elbD* homolog) containing fatty acyl CoA–like reductase, acyl-CoA synthetase, thiolation and acylglycerolphosphate acyltransferase domains. The NRPS-like core gene was consistently flanked by hydrolases/dehydratases, as well as oxidoreductases and genes involved in fatty acid biosynthesis (Fig. 5). These GCFs show similarity to a known BGC encoding the production of vinyl-/alkyl-ether lipids (VEPE/AEPE) (MiBIG accession: BGC0000871). The VEPE/AEPE lipids are produced by *Myxococcus xanthus* DK 1622 as extracellular signals guiding fruiting body morphogenesis and sporulation (44,73). It is postulated that these lipids are generated via modification of phospholipids originating from the cell membrane with participation of BGC0000871’s genes *elbB, D & E*, as well as an additional desaturase *CarF* (73,74). An *elbD* homolog of poribacterial origin has also been identified by Lorenzen et. al (44).

**Figure 5.**
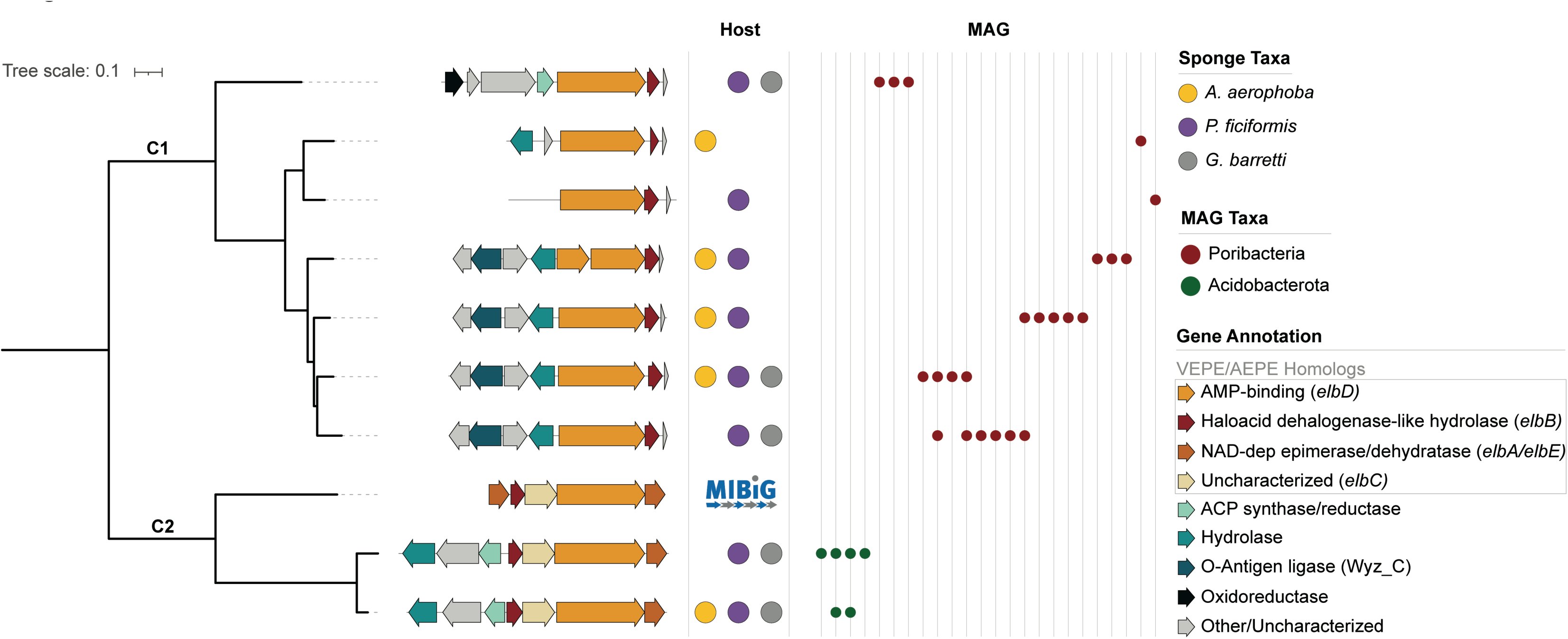
Distribution and characterization of the VEPE related GCFs. FastTree phylogeny of VEPE-associated GCFs, each characterized by a representative BGC, based on sequence similarity of the A-domain (AMP-binding). Detailed overview of these GCFs in Supplement 5.2 C1 and C2 refer to Clade 1 and 2. Genes are coloured based on predicted function. Each BGC is annotated by their presence in host sponge species and in MAGs.

We observe architectural changes in the adjacent genes of the cluster across the set that are congruent with the clades generated based on sequence similarity of the A-domain (Fig. 5). Additional analysis of adenylation domain active site specificity-conferring residues (AdenylPred (75), Supplement 5.1) indicates potentially different functional classes and substrate specificities for the two clades, which suggests diversity in the chemical compounds produced by the encoded machinery. Gene clusters from the two clades are associated with distinct bacterial taxa, being specific to Acidobacterota and Poribacteria (GTDB r95) MAGs, with any given MAG harbouring a maximum of 2 of the 9 GCFs (Supplement 5.3).

Ether lipids in sponges are often linked to pathogen defense by showing antimicrobial activity (76–79). However, the biosynthetic origin and pathway for these molecules is currently undescribed. Ultimately experimental work will be necessary to determine the function and biosynthesis of these ether lipids within the sponge holobiont.

### High RiPP diversity in sponge bacterial symbionts

Even though sponges are recognized as extensive sources of diverse NPs, their inventory of ribosomally synthesized and post-translationally modified peptides (RiPP) has remained largely undescribed, with one exception being the proteusin polytheonamides (15,23,80). Here we expand the known repertoire in sponges and showcase sponges’ RiPP diversity by identifying 17 uncharacterized RiPP families which seem to be widespread in sponge holobionts. In some of these cases, GCFs are present across all sponge species, which points towards an important role of the produced NPs in the context of sponge-microbe symbiosis (Fig. 6).

**Figure 6.**
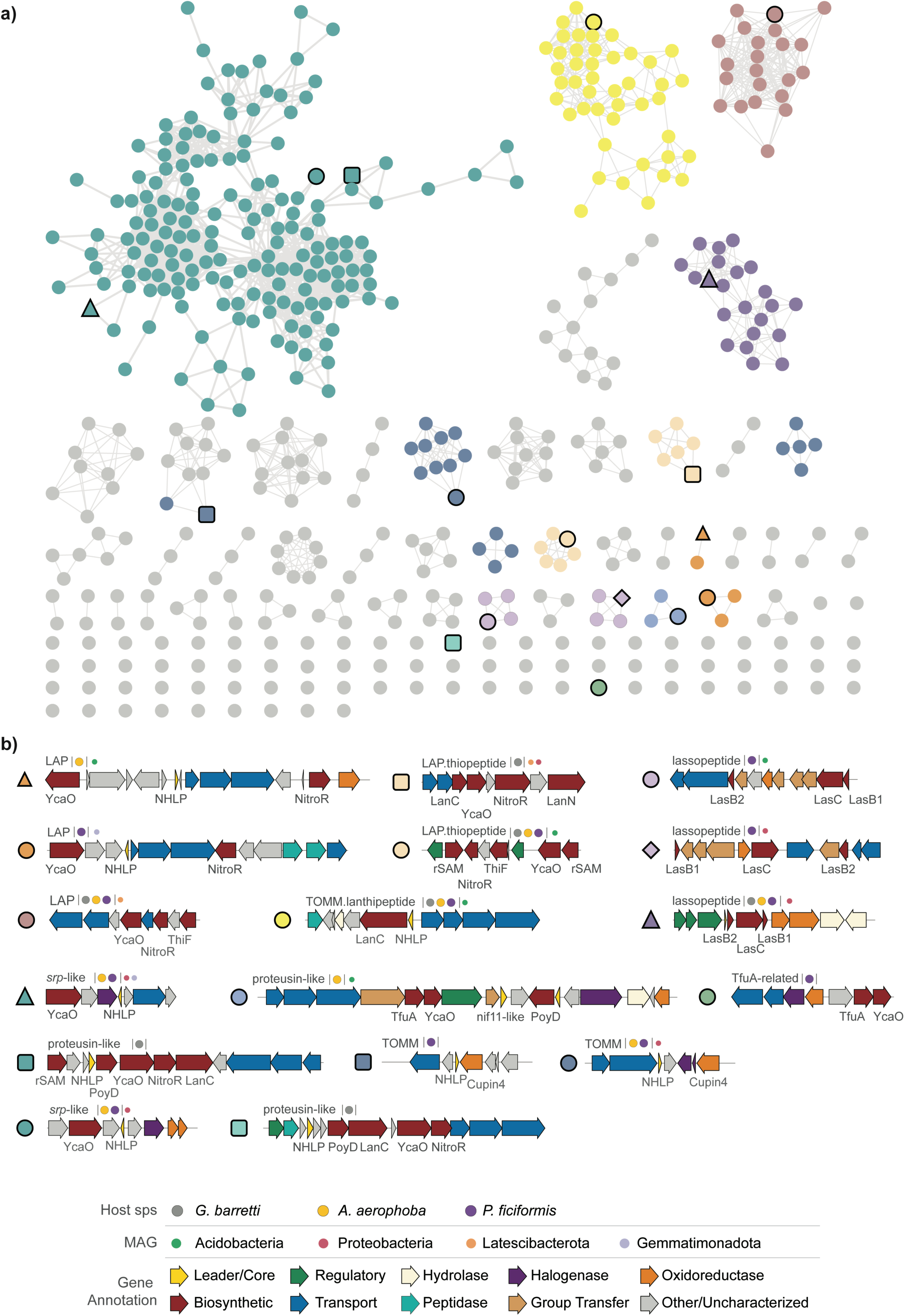
An overview of RiPP diversity. **a)** RiPP BGCs represented in a BiG-SCAPE generated similarity network. Nodes are coloured by biosynthetic class/characteristic, **b)** Representative BGCs from each GCF are depicted and characterized by predicted class, presence in a host sponge and MAG (GTDB r95). Genes are coloured by predicted function.

RiPPs constitute a class of natural products that is produced from a ribosomally synthesized precursor peptide, composed of an N-terminal leader and a C-terminal core, which is modified by biosynthetic enzymes encoded in the BGC, and matured in a final proteolytic cleavage step (81). We see a high incidence of RiPP BGCs with predicted nitrile hydratase-like leader peptide (NHLP) domains, which have been identified as proteusin precursor peptides, as well as nif11-like leader peptide domains associated with thiazole/oxazole-modified microcins (TOMM) precursor peptides (82). Encoded within these BGCs are a number of modifying enzymes that span several RiPP subclasses: YcaO cyclodehydratases (83,84), radical S-adenosylmethionine (rSAM) enzymes (85) including PoyD (80), lanthionine synthetase C-like (LanC) enzymes (86), nitroreductases (NitroR) (87) and TfuA-like enzymes (88). Other modifying enzymes include oxidoreductases, group transferases and halogenases. Several RiPP subclasses contain YcaO-generated azol(in)e heterocycles (84), such as the linear azol(in)e-containing peptides (LAP) and thiopeptides, and are often linked to pharmacologically interesting bioactivities (89–91). The potential for azol(in)e-containing RiPPs has been found widespread in bacterial genomes (92), but has only recently been reported in marine sponge microbiomes with the identification of the *srp* clusters by Nguyen *et*.*al* (15). We predict an additional structurally diversified set of azol(in)e-containing RiPPs to this repertoire, expanding the combinatorial space of modifying enzymes and hence the potential for production of bioactive NPs.

Additionally we observe four GFCs that harbour a cupin-4 modifying enzyme in addition to a NHLP domain, which has also been connected to proteusin biosynthesis (93). The cupin superfamily is functionally highly diverse, with reported hydroxylation, epoxidation, dioxygenation, decarboxylation, dehydration, and halogenation activities (94–96). Although these genes appear in marine environments, they are not yet regarded as a typical constituent of the sponge microbiome biosynthetic array (97,98). We further identified several GCFs predicted to be responsible for the production of highly modified lassopeptides, encoding for additional hydrolases, oxidoreductases, and group transferases (sulfo-, glycol-, nucletotidyl-). These include GCFs that are shared among all sponge species. Lassopeptides have been found in some sponge-associated microbiomes (99,100), but their presence together with diverse flanking enzyme-encoding genes has not been previously reported in this environmental niche. Whilst the functions of these BGCs and their metabolic products remains elusive, this dataset expands and diversifies the set of sponge-holobiont-encoded RiPPs, and predicts sponge microbiomes to be a prolific sources of RiPP NPs.

## Conclusion

Here we describe a systematic analysis of the secondary metabolite diversity landscape of several marine sponge holobionts, whose metagenomes contain a large number of uncharacterized BGCs. We show that while there is high biosynthetic diversity that is unique to each sponge species, a small functional core is conserved throughout the studied sponge species. This functional core includes a novel group of NRPS-like ether lipid-associated GCFs widespread across sponges and a number of RiPPs. We recovered a consistent set of biosynthetically potent MAGs from the metagenomes, with Acidobacteriota and Latescibacterota standing out as potential prolific NP producers. This study thus contributes to a growing body of research uncovering the network of NPs with elusive functions that are generated by marine sponge microbiomes.

## Acknowledgments

The authors wish to thank Ellen Kenchington for providing the Canadian *Geodia barretti* samples, the late Hans Tore Rapp for his help in sampling the Norwegian *G. barretti* samples, and dr. Vasilis Gerovasileiou and HCMR (Hellenic Centre for Marine Research) for the collaboration in sampling the Petrosia ficiformis samples. We thank Vittorio Tracanna, Joris Louwen, Michelle Schorn for valuable advice and discussions. This research was financially supported by the VLAG NWO PhD project “*Mare incognita*” to CL, the European Commission through Horizon2020 project SponGES (Grant agreement ID: 679849) to DS and AG (Asimenia) and the BluePharmTrain project (Grant agreement ID: 607786) to DS.

## Author Contributions

CL: Research design. *G. barretti* and Seawater_ATL eDNA extraction. Seawater, *G. barretti, A. aerophoba* sequence QC-filtering and assembly. All samples BGC prediction and characterization, binning, taxonomy profile prediction, Nonpareil estimates and downstream analysis. Writing the manuscript.

A.G (Anastasia): *P. ficiformis* sampling, eDNA extraction, QC-filtering and assembly

A.G (Asimenia): *G. barretti* and Seawater_ATL sampling and eDNA extraction

M.C.dM: SW_Med and *A. aerophoba* sampling and eDNA extraction

J.vdO: Research design, manuscript reviewing/editing

M.H.M: Research design, manuscript reviewing/editing

D.S: Research design, manuscript reviewing/editing, *G. barretti* and Seawater_ATL sampling

## Data availability

The data for this study have been deposited in the European Nucleotide Archive (ENA) at EMBL-EBI under accession number PRJEB51534. Python scripts created for this analysis are available at https://github.com/CatarinaCarolina/sponge_meta_BGC.

## Supplementary Material

- S1: Sample Metadata
- S2.1: SUP and swf GCF
- S2.2: SUP and swf MAG distribution
- S3: Diversity statistical tests
- S4: iTOL with taxa annotation
- S5.1: Ether lipid AdenylPred predictions
- S5.2: Ether lipid GCFs
- S5.3: Ether lipid GCFs in MAGs

