## Supplementary Material for "Comparative metagenomic analysis of biosynthetic diversity across sponge microbiomes highlights metabolic novelty, conservation and diversification"

This PDF file includes:

S1: Sample Metadata
S2.1: SUP and swf GCFs
S2.2: SUP and swf MAG distribution
S3: Diversity statistic test
S4: MAG iTOL table with taxonomy annotation
S5.1: AdenylPred predictions
S5.2: Ether lipid GCFs
S5.3: Ether lipid MAG distribution

**Supplement 1. Sample metadata**

| Name | Type | Species | Depth_avg (m) | Location | Coordinates | Country | Sampling Year | Sequencing Year | Sequencing technology |
|---|---|---|---|---|---|---|---|---|---|
| Aply16 | Sponge Tissue | Aplysina aerophoba | 10,25 | Cala Montgo | N42°06'52.20", E3°10'06.52" | Spain | 2014 | 2014 | Hybrid Illumina Pacbio |
| Aply21 | Sponge Tissue | Aplysina aerophoba | 10,25 | Cala Montgo | N42°06'52.20", E3°10'06.52" | Spain | 2014 | 2014 | Hybrid Illumina Pacbio |
| Aply22 | Sponge Tissue | Aplysina aerophoba | 10,25 | Cala Montgo | N42°06'52.20", E3°10'06.52" | Spain | 2014 | 2014 | Hybrid Illumina Pacbio |
| Aply23 | Sponge Tissue | Aplysina aerophoba | 10,25 | Cala Montgo | N42°06'52.20", E3°10'06.52" | Spain | 2014 | 2014 | Hybrid Illumina Pacbio |
| Pf4 | Sponge Tissue | Petrocia ficiformis | 5,5 | Sfakia, Crete (Semi-dark zone) | N35°12'0.7", E24°7'09.8" | Greece | 2018 | 2018 | Illumina |
| Pf5 | Sponge Tissue | Petrocia ficiformis | 5,5 | Sfakia, Crete (Semi-dark zone) | N35°12'0.7", E24°7'09.8" | Greece | 2018 | 2018 | Illumina |
| Pf6 | Sponge Tissue | Petrocia ficiformis | 5,5 | Sfakia, Crete (Semi-dark zone) | N35°12'0.7", E24°7'09.8" | Greece | 2018 | 2018 | Illumina |
| Pf7 | Sponge Tissue | Petrocia ficiformis | 5,5 | Sfakia, Crete (Dark zone) | N35°12'0.7", E24°7'09.8" | Greece | 2018 | 2018 | Illumina |
| Pf8 | Sponge Tissue | Petrocia ficiformis | 5,5 | Sfakia, Crete (Dark zone) | N35°12'0.7", E24°7'09.8" | Greece | 2018 | 2018 | Illumina |
| Pf9 | Sponge Tissue | Petrocia ficiformis | 5,5 | Sfakia, Crete (Dark zone) | N35°12'0.7", E24°7'09.8" | Greece | 2018 | 2018 | Illumina |
| Pf10 | Sponge Tissue | Petrocia ficiformis | 5,5 | Sfakia, Crete (Entrance) | N35°12'0.7", E24°7'09.8" | Greece | 2018 | 2018 | Illumina |
| Pf11 | Sponge Tissue | Petrocia ficiformis | 5,5 | Sfakia, Crete (Entrance) | N35°12'0.7", E24°7'09.8" | Greece | 2018 | 2018 | Illumina |
| Pf12 | Sponge Tissue | Petrocia ficiformis | 5,5 | Sfakia, Crete (Entrance) | N35°12'0.7", E24°7'09.8" | Greece | 2018 | 2018 | Illumina |
| gb1 | Sponge Tissue | Geodia barretti | 450 | Scengsbukt-Korsfjord | N60°8'8", E5°6'42" | Norway | 2017 | 2018 | Illumina |
| gb2_2 | Sponge Tissue | Geodia barretti | 150 | Scengsbukt-Korsfjord | N60°8'8", E5°6'42" | Norway | 2017 | 2018 | Illumina |
| gb4_2 | Sponge Tissue | Geodia barretti | 150 | Scengsbukt-Korsfjord | N60°8'8", E5°6'42" | Norway | 2017 | 2018 | Illumina |
| gb5_2 | Sponge Tissue | Geodia barretti | 150 | Scengsbukt-Korsfjord | N60°8'8", E5°6'42" | Norway | 2017 | 2018 | Illumina |
| gb6 | Sponge Tissue | Geodia barretti | 150 | Scengsbukt-Korsfjord | N60°8'8", E5°6'42" | Norway | 2017 | 2018 | Illumina |
| gb7 | Sponge Tissue | Geodia barretti | 450 | Scengsbukt-Korsfjord | N60°8'8", E5°6'42" | Norway | 2017 | 2018 | Illumina |
| gb8_2 | Sponge Tissue | Geodia barretti | 450 | Scengsbukt-Korsfjord | N60°8'8", E5°6'42" | Norway | 2017 | 2018 | Illumina |
| gb9 | Sponge Tissue | Geodia barretti | 450 | Scengsbukt-Korsfjord | N60°8'8", E5°6'42" | Norway | 2017 | 2018 | Illumina |
| gb10 | Sponge Tissue | Geodia barretti | 450 | Scengsbukt-Korsfjord | N60°8'8", E5°6'42" | Norway | 2017 | 2018 | Illumina |
| gb126 | Sponge Tissue | Geodia barretti | 1213 | Davis Strait | N62°52'15.1", W58°37'34.32" | Canada | 2015 | 2018 | Illumina |
| gb278 | Sponge Tissue | Geodia barretti | 1335 | Davis Strait | N61°53'36.13", W60°7'57.612" | Canada | 2014 | 2018 | Illumina |
| gb305 | Sponge Tissue | Geodia barretti | 1437 | Davis Strait | N62°31'6.24", W59°58'13.872" | Canada | 2014 | 2018 | Illumina |
| gb1_f | Filtered Sea Water | N.a. Seawater Atl | 150 | Scengsbukt-Korsfjord | N60°8'8", E5°6'42" | Norway | 2017 | 2018 | Illumina |
| gb2_f | Filtered Sea Water | N.a. Seawater Atl | 150 | Scengsbukt-Korsfjord | N60°8'8", E5°6'42" | Norway | 2017 | 2018 | Illumina |
| gb3_f | Filtered Sea Water | N.a. Seawater Atl | 150 | Scengsbukt-Korsfjord | N60°8'8", E5°6'42" | Norway | 2017 | 2018 | Illumina |
| gb5_6_f | Filtered Sea Water | N.a. Seawater Atl | 450 | Scengsbukt-Korsfjord | N60°8'8", E5°6'42" | Norway | 2017 | 2018 | Illumina |
| gb9_f | Filtered Sea Water | N.a. Seawater Atl | 450 | Scengsbukt-Korsfjord | N60°8'8", E5°6'42" | Norway | 2017 | 2018 | Illumina |
| gb10_f | Filtered Sea Water | N.a. Seawater Atl | 450 | Scengsbukt-Korsfjord | N60°8'8", E5°6'42" | Norway | 2017 | 2018 | Illumina |
| sw_7 | Filtered Sea Water | N.a. Seawater Med | 10,25 | Cala Montgo | N42°06'52.20", E3°10'06.52" | Spain | 2014 | 2014 | Illumina |
| sw_8 | Filtered Sea Water | N.a. Seawater Med | 10,25 | Cala Montgo | N42°06'52.20", E3°10'06.52" | Spain | 2014 | 2014 | Illumina |
| sw_9 | Filtered Sea Water | N.a. Seawater Med | 10,25 | Cala Montgo | N42°06'52.20", E3°10'06.52" | Spain | 2014 | 2014 | Illumina |

**Supplement 2.1. SUP-like and swf-like GCFs. a) SUP and swf related clans in BiG-SCAPE PKSI network. b) GCF composition of SUP and swf-like GCF examples.**

**a)**

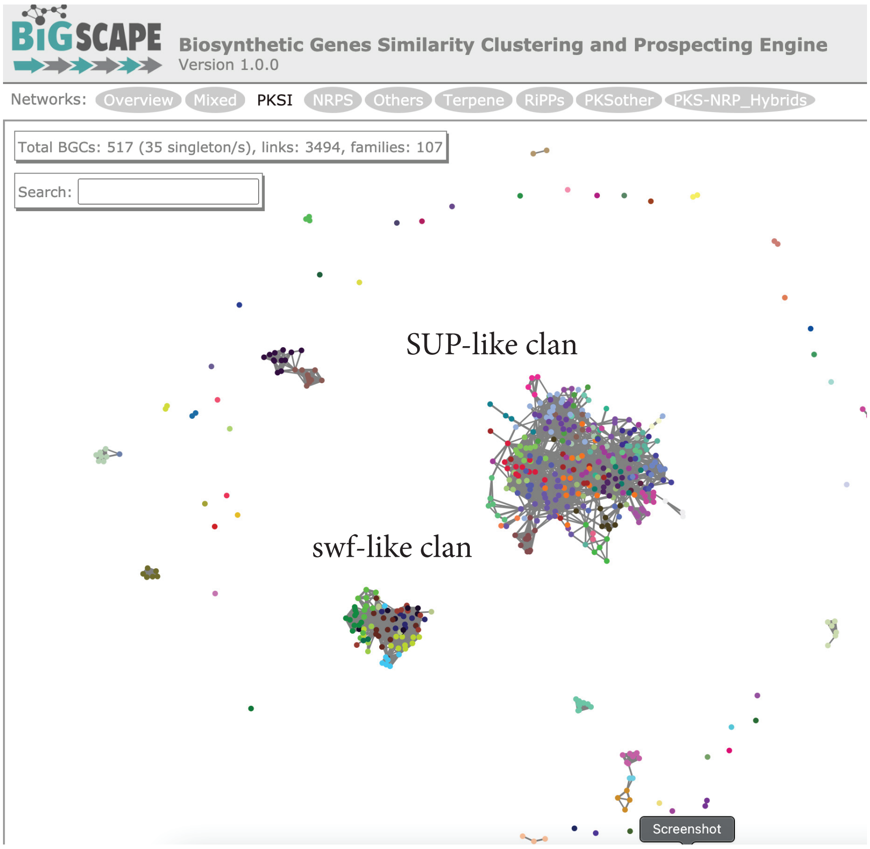

**b**

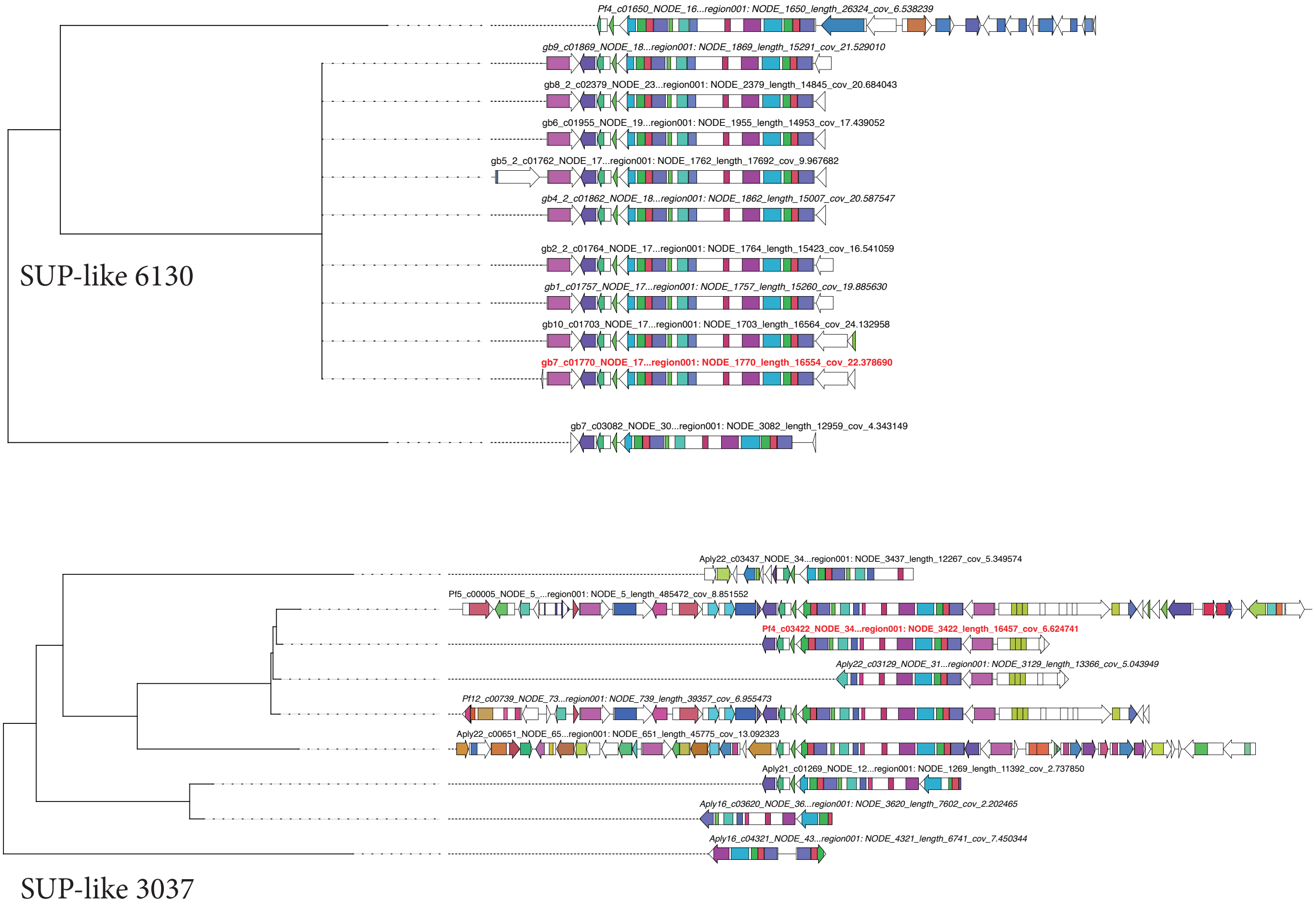

### swf-like 2399

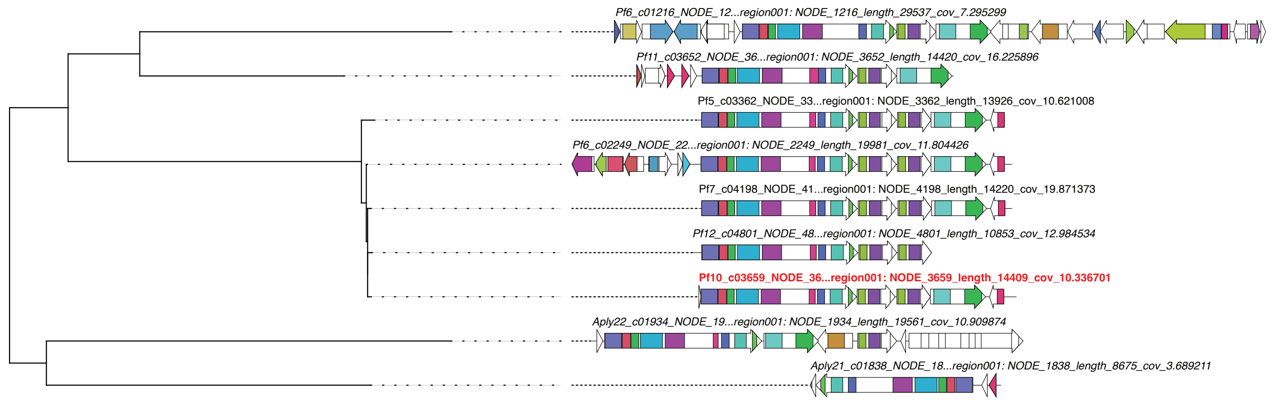

### swf-like 6055

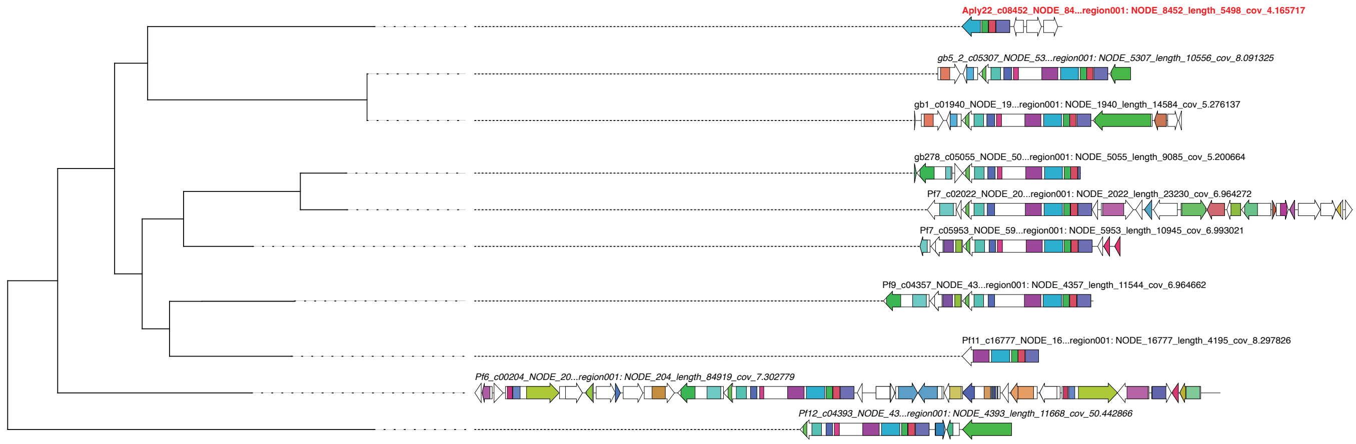

**Supplement 2.2. SUP-like and swf-like GCF examples binned in MAGs, characterised by MAG distribution and phylum taxonomy**

| **SUP-like example** | | |
|---|---|---|
| GCF | Sponge species | |
| **6130** | **Aplysina,Petrosia,Geodia** | |
| Encoding bin | Bin presence host | Bin taxonomy |
| Pf4_bin.62.fa | Petrosia | p__Chloroflexota |
| Pf7_bin.11.fa | Petrosia | p__Spirochaetota |
| Pf9_bin.9.fa | Petrosia | p__Proteobacteria |
| Pf8_bin.5.fa | Petrosia | p__Proteobacteria |
| Pf6_bin.10.fa | Petrosia | p__Proteobacteria |
| Pf5_bin.39.fa | Petrosia | p__Acidobacteriota |
| Pf8_bin.46.fa | Aplysina,Petrosia | p__Acidobacteriota |
| gb5_2_bin.68.fa | Geodia | p__Acidobacteriota |

| **swf-like example** | | |
|---|---|---|
| GCF | Sponge species | |
| **2399** | **Aplysina, Petrosia** | |
| Encoding bin | Bin presence in host | Bin taxonomy |
| Aply22_bin.37.fa | Aplysina | p__Nitrospirota |
| Pf7_bin.67.fa | Aplysina, Petrosia | p__Latescibacterota |
| Pf10_bin.11.fa | Petrosia | p__Acidobacteriota |

| **swf-like example** | | |
|---|---|---|
| GCF | Sponge species | |
| **6055** | **Aplysina,Petrosia,Geodia** | |
| Encoding bin | Bin presence in host | Bin taxonomy |
| gb10_bin.62.fa | Geodia | p__Latescibacterota |

**Supplement 5.2. Ether Lipid A domain AdenylPred predictions, C1 and C2 refer to clades 1 and 2 as in main Fig.5. Prediction probabilities are averaged within each clade.**

| Query name | Predicted functional class (FC) | FC prediction probability | Predicted substrate specificity (SS) | SS prediction probability |
|---|---|---|---|---|
| **C1 AMP_binding** | Aryl-CoA ligase | **0.48** | cinnamate and succinylbenzoate derivatives | **0.34** |
| **C2 AMP_binding** | Long chain acyl-CoA synthetase | **0.54** | C13 through C17 | **0.45** |
| **BGC0000871_MXAN_1528_AMP-binding** | Long chain acyl-CoA synthetase | **0.67** | C13 through C17 | **0.6** |

#### Supplement 3. Alpha and beta diversity statistical tests

##### PERMANOVA results

```
Input    phyloFlash_compare.6.ntu_table.braycurtis
method name                PERMANOVA
test statistic name         pseudo-F
sample size                       35
number of groups                   6
test statistic               55.7199
p-value                        0.001
number of permutations           999

input    BiG_MAP.all_RPKM_NORM.braycurtis
method name                PERMANOVA
test statistic name         pseudo-F
sample size                       35
number of groups                   6
test statistic               97.4049
p-value                        0.001
number of permutations           999
```

##### Shannon alpha diversity based tests

```
Sponge Mean BGC: 8.45
Sponge STD BGC: 0.13
Sponge VAR BGC: 0.02
Sponge Mean Taxa: 5.99
Sponge STD Taxa: 0.26
Sponge VAR Taxa: 0.07
geodianor Mean BGC: 8.36
geodianor STD BGC: 0.07
geodianor VAR BGC: 0.01
geodianor Mean Taxa: 5.71
geodianor STD Taxa: 0.08
geodianor VAR Taxa: 0.01
geodiacan Mean BGC: 8.29
geodiacan STD BGC: 0.10
geodiacan VAR BGC: 0.01
geodiacan Mean Taxa: 6.13
geodiacan STD Taxa: 0.11
geodiacan VAR Taxa: 0.01
petrosia Mean BGC: 8.59
petrosia STD BGC: 0.05
petrosia VAR BGC: 0.01
petrosia Mean Taxa: 6.09
petrosia STD Taxa: 0.10
petrosia VAR Taxa: 0.01
aplysina Mean BGC: 8.46
aplysina STD BGC: 0.08
aplysina VAR BGC: 0.01
aplysina Mean Taxa: 6.36
aplysina STD Taxa: 0.17
aplysina VAR Taxa: 0.03
Water Mean BGC: 6.94
Water STD BGC: 0.74
Water VAR BGC: 0.56
Water Mean Taxa: 6.96
Water STD Taxa: 0.16
Water VAR Taxa: 0.03
```

```
kruskal BGC all sponge:  KruskalResult(statistic=17.10, pvalue=0.0007)
kruskal BGC gbs:  KruskalResult(statistic=1.029, pvalue=0.31)
kruskal BGC gb aa:  KruskalResult(statistic=3.92, pvalue=0.048)
kruskal BGC pf aa:  KruskalResult(statistic=4.67, pvalue=0.031)
kruskal BGC gb pf : KruskalResult(statistic=12.33, pvalue=0.0005)
kruskal taxa all:  KruskalResult(statistic=19.97, pvalue=0.0002)
kruskal taxa gbs:  KruskalResult(statistic=6.43, pvalue=0.011)
kruskal taxa gb aa:  KruskalResult(statistic=8.0, pvalue=0.005)
kruskal taxa pf aa:  KruskalResult(statistic=5.36, pvalue=0.021)
kruskal taxa gb pf : KruskalResult(statistic=13.5, pvalue=0.0002)

All Coefficient of Determination -18.64
All Spearman -0.4159663865546219 0.013
```

**Supplement 4. MAG iTOL tree with bin name and taxonomy annotations.**

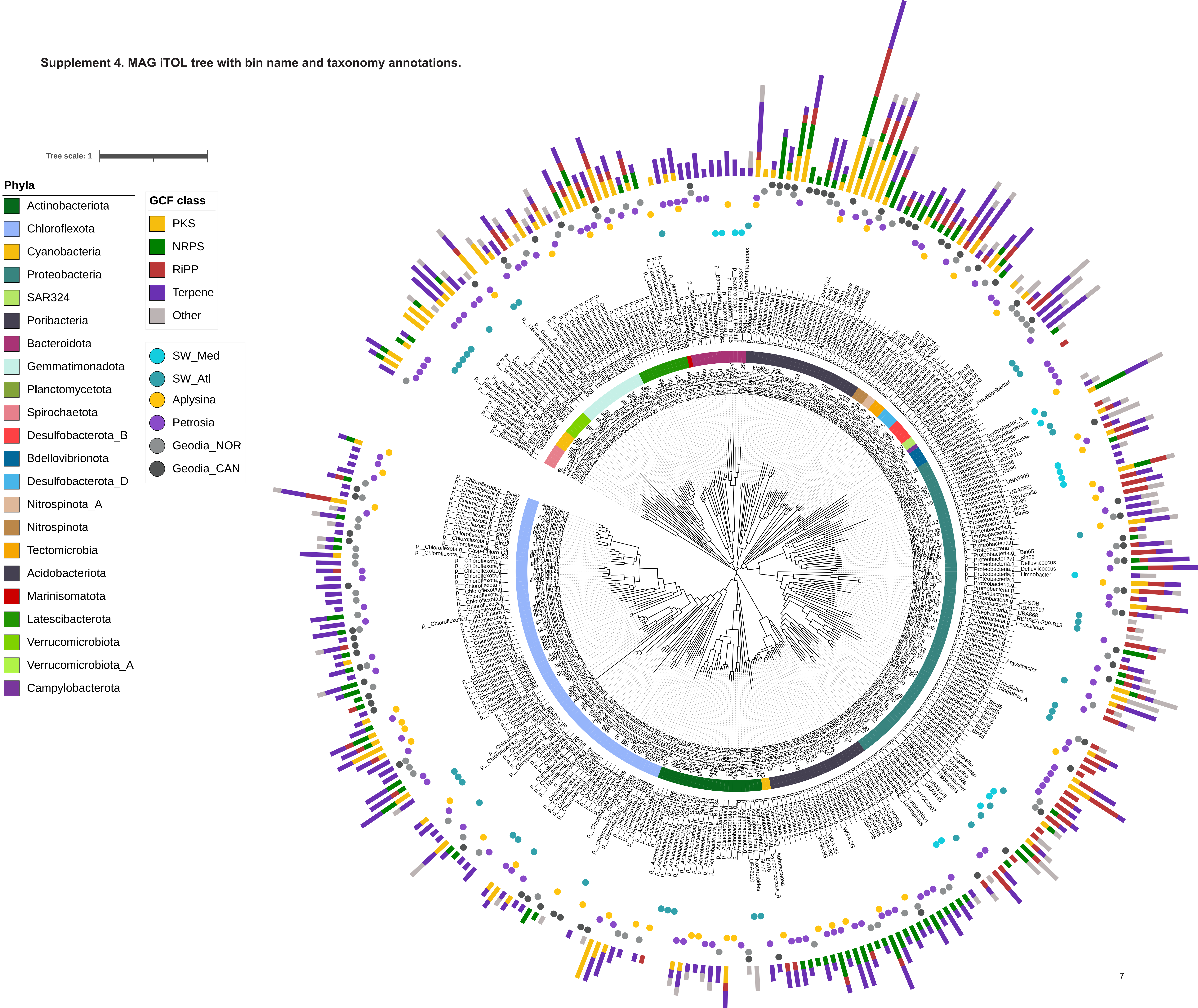

**Supplement 5.2. Detailed depiction of all GCFs captured by CORASON and included in ether lipid search. GCfs are numbered and red box indicates chosen representative.**

5496

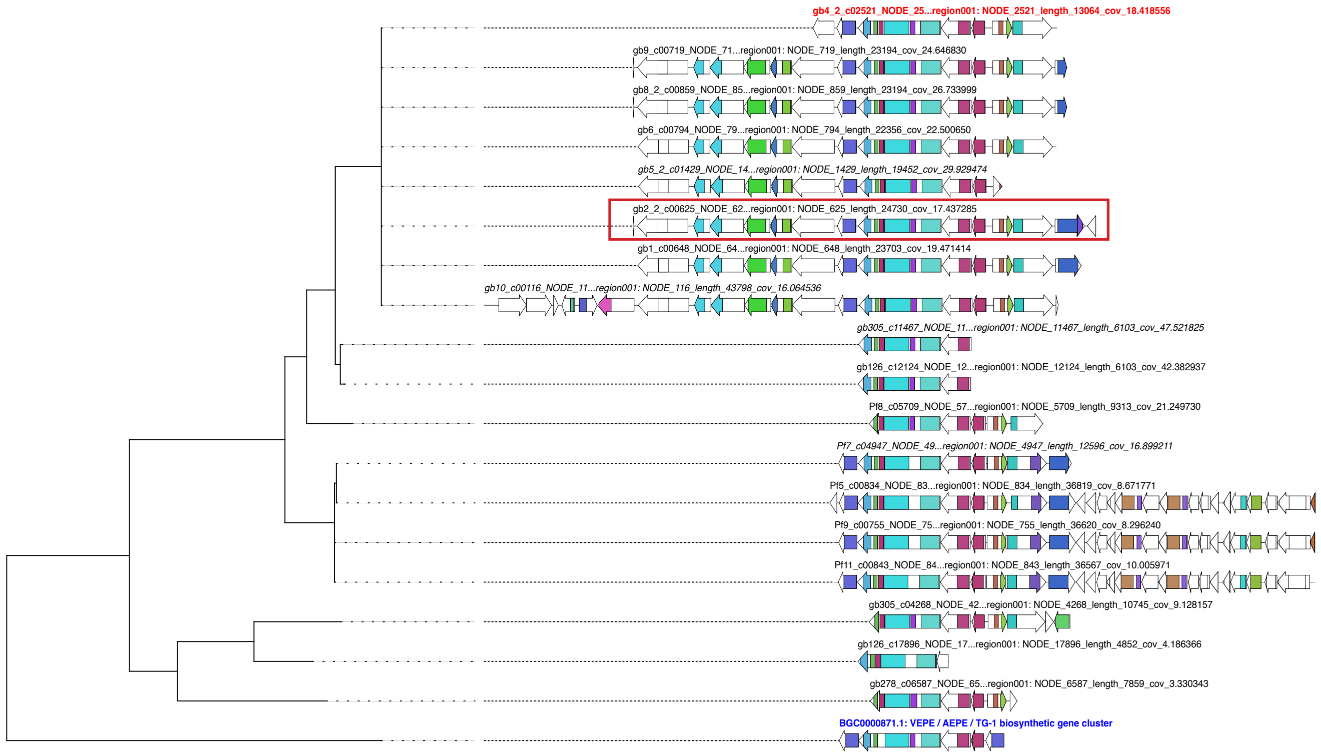

380

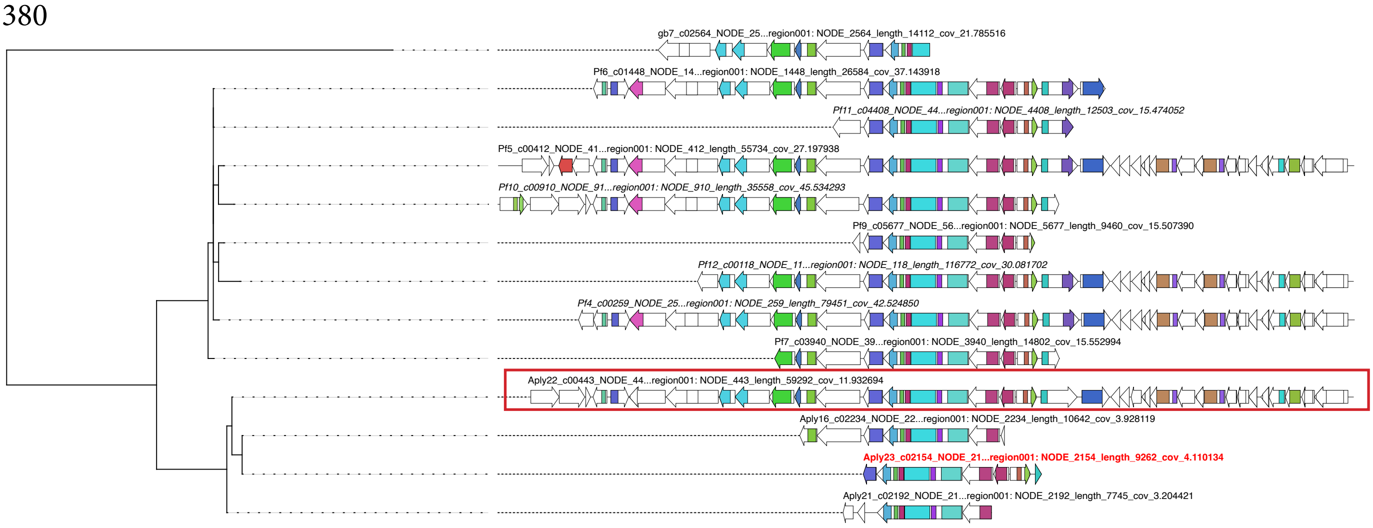

3574

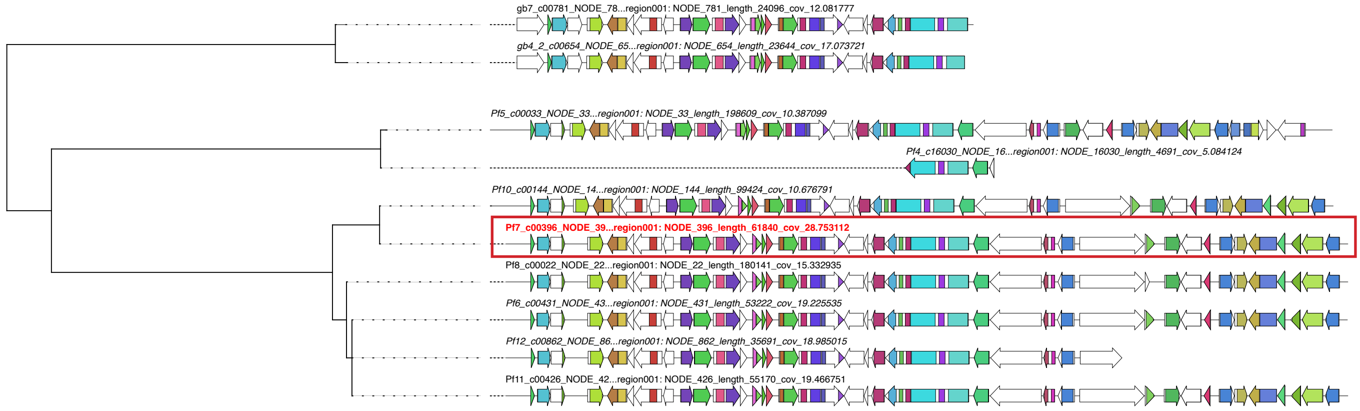

6120

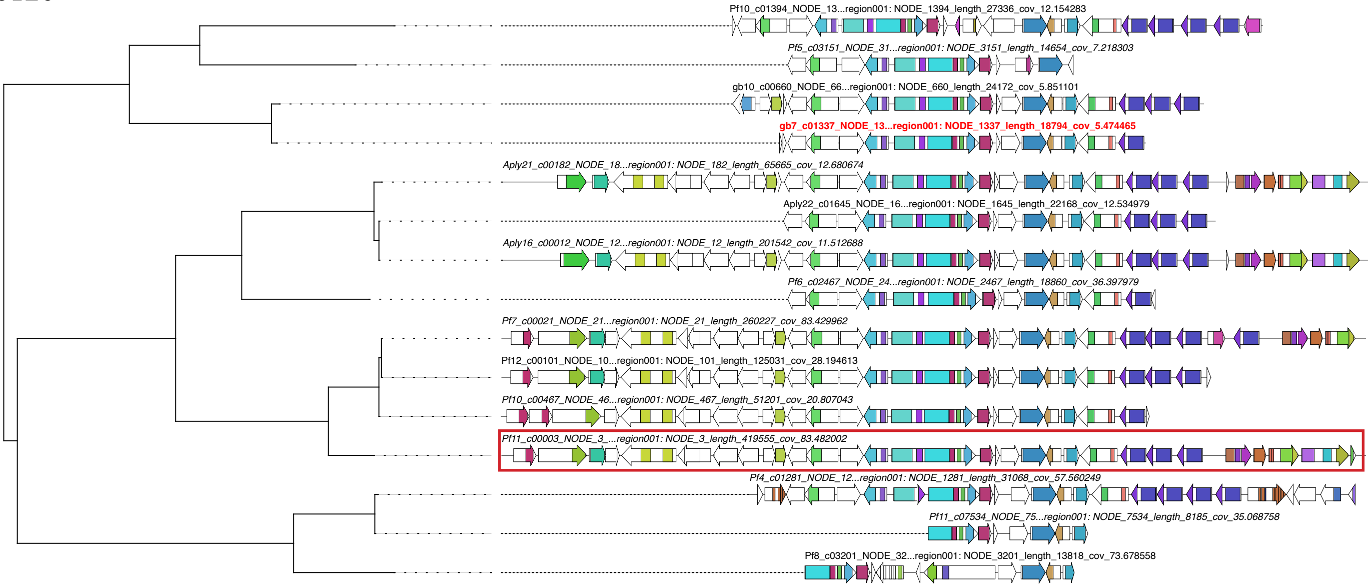

4988

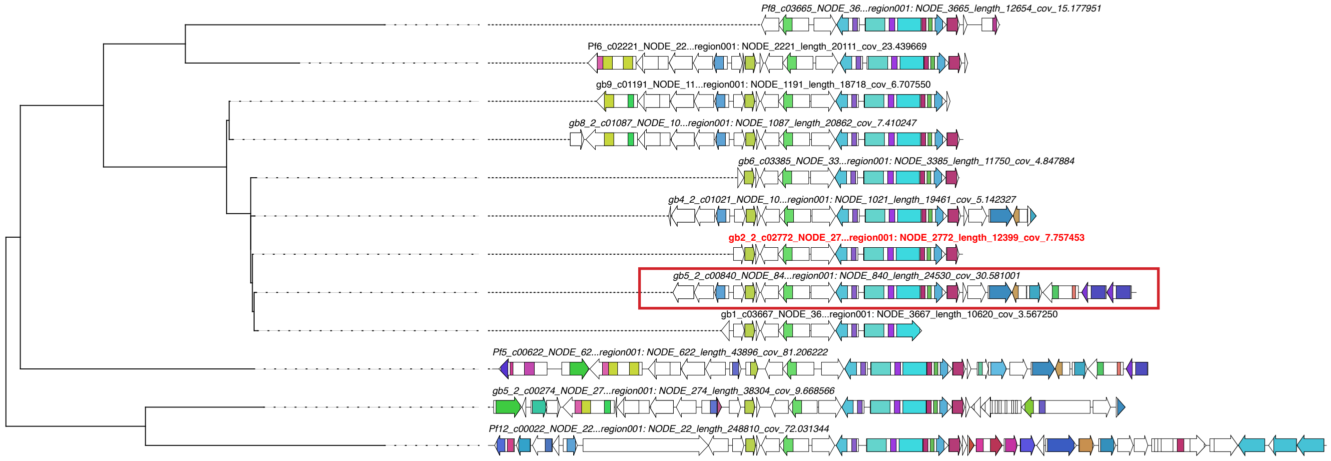

3047

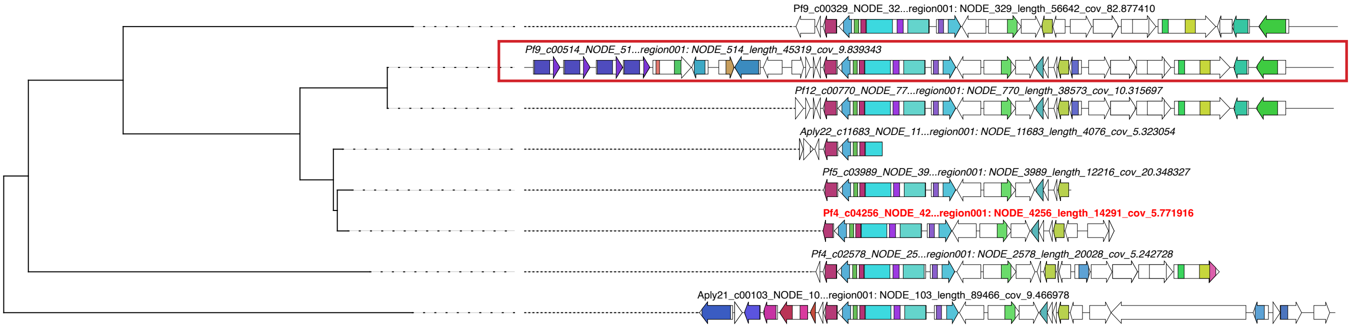

4093

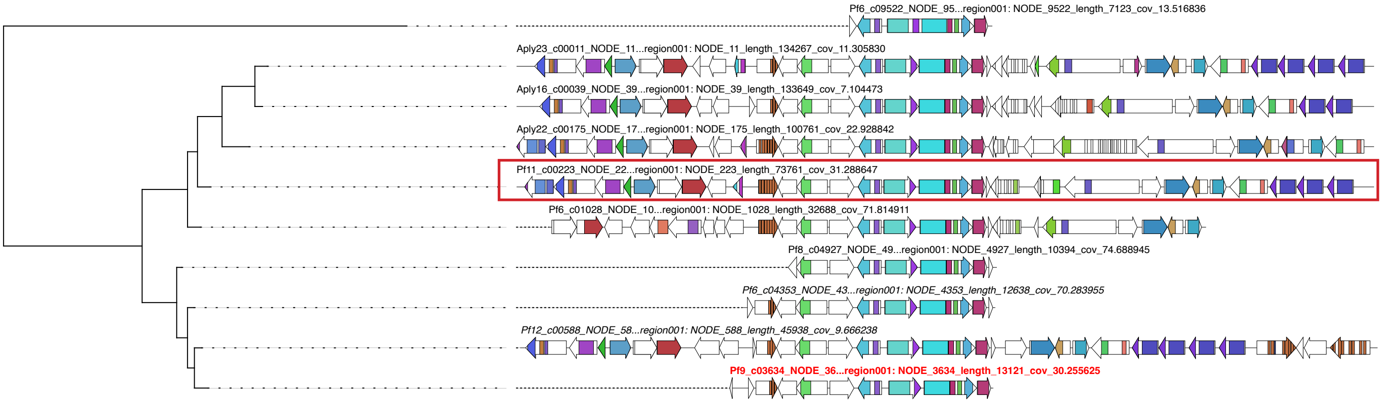

172

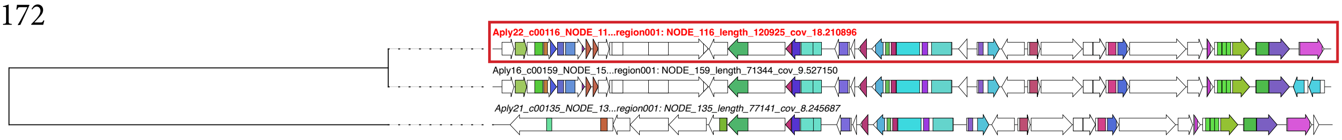
Aply22_c00116_NODE_11...region001: NODE_116_length_120925_cov_18.210896
Aply16_c00159_NODE_15...region001: NODE_159_length_71344_cov_9.527150
Aply21_c00135_NODE_13...region001: NODE_135_length_77141_cov_8.245687

4034

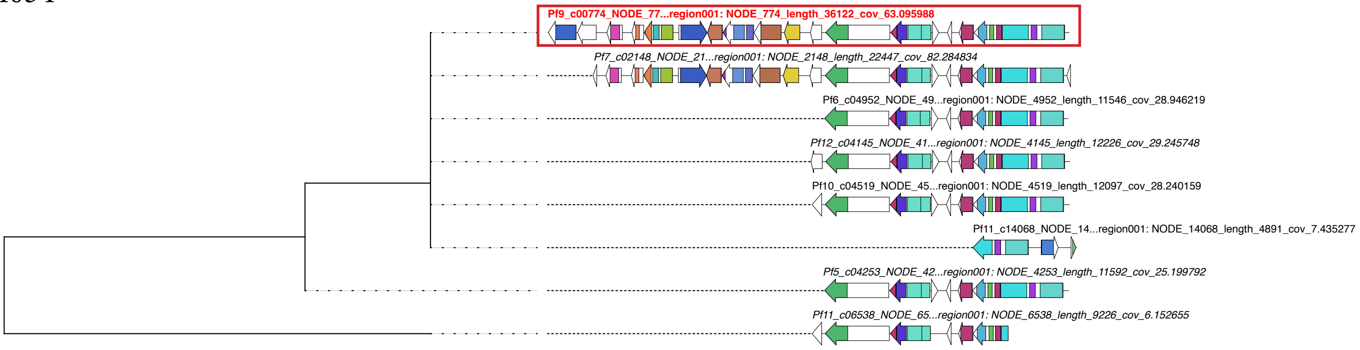
Pf9_c00774_NODE_77...region001: NODE_774_length_36122_cov_63.095988
Pf7_c02148_NODE_21...region001: NODE_2148_length_22447_cov_82.284834
Pf6_c04952_NODE_49...region001: NODE_4952_length_11546_cov_28.946219
Pf12_c04145_NODE_41...region001: NODE_4145_length_12226_cov_29.245748
Pf10_c04519_NODE_45...region001: NODE_4519_length_12097_cov_28.240159
Pf11_c14068_NODE_14...region001: NODE_14068_length_4891_cov_7.435277
Pf5_c04253_NODE_42...region001: NODE_4253_length_11592_cov_25.199792
Pf11_c06538_NODE_65...region001: NODE_6538_length_9226_cov_6.152655

#### Supplement 5.3. MAGs containing ether lipid GCFs, and characterized by phylum taxonomy

| Taxa | GCF/MAG | 5496 | 380 | 3574 | 6120 | 4988 | 3047 | 4093 | 172 | 4034 |
|---|---|---|---|---|---|---|---|---|---|---|
| Acidobacteria | Pf5_bin.39 | x | | | | | | | | |
| | Pf4_bin.17 | x | x | | | | | | | |
| | gb6_bin.52 | x | x | | | | | | | |
| | gb126_bin.28 | x | | | | | | | | |
| Poribacteria | Pf8_bin.6 | | | x | | | | | | |
| | Pf5_bin.13 | | | x | | | | | | |
| | Aply16_bin.2 | | | | x | | | | | |
| | Pf10_bin.3 | | | | x | x | | | | |
| | Pf11_bin.65 | | | | x | | | | | |
| | gb4_2_bin.50 | | | | x | x | | | | |
| | Pf12_bin.33 | | | | | x | x | | | |
| | Pf5_bin.4 | | | | | x | | | | |
| | Pf8_bin.49 | | | | | x | | | | |
| | gb5_2_bin.34 | | | | | x | | | | |
| | Pf5_bin.37 | | | | | | x | | | |
| | Pf9_bin.14 | | | | | | x | | | |
| | Pf9_bin.10 | | | | | | x | | | |
| | Aply23_bin.19 | | | | | | | x | | |
| | Pf5_bin.52 | | | | | | | x | | |
| | Pf9_bin.24 | | | | | | | x | | |
| | Aply22_bin.11 | | | | | | | | x | |
| | Pf5_bin.31 | | | | | | | | | x |
